# A comprehensive model of DNA fragmentation for the preservation of High Molecular Weight DNA

**DOI:** 10.1101/254276

**Authors:** Tomas Klingström, Erik Bongcam-Rudloff, Olga Vinnere Pettersson

## Abstract

For long-read sequencing applications, shearing of DNA is a significant issue as it limits the read-lengths generated by sequencing. During extraction and storage of DNA the DNA polymers are susceptible to physical and chemical shearing. In particular, the mechanisms of physical shearing are poorly understood in most laboratories as they are of little relevance to commonly used short-read sequencing technologies. This study draws upon lessons learned in a diverse set of research fields to create a comprehensive theoretical framework for obtaining high molecular weight DNA (HMW-DNA) to support improved quality management in laboratories and biobanks for long-read sequencing applications.

Under common laboratory conditions physical and chemical shearing yields DNA fragments of 5-35 kilobases (kb) in length. This fragment length is sufficient for DNA sequencing using short-read technologies but for Nanopore sequencing, linked reads and single molecular real time sequencing (SMRT) poorly preserved DNA will limit the length of the reads generated.

The shearing process can be divided into physical and chemical shearing which generates different patterns of fragmentation. Exposure to physical shearing creates a characteristic fragment length where the main cause of shearing is shear stress induced by turbulence. The characteristic fragment length is several thousand base pairs longer than the reads produced by short-read sequencing as the shear stress imposed on short DNA fragments is insufficient to shear the DNA. This characteristic length can be measured using gel electrophoresis or instruments for DNA fragment analysis,. Chemical shearing generates randomly distributed fragment lengths visible as a smear of DNA below the peak fragment length. By measuring the peak of the DNA fragment length distribution and the proportion of very short DNA fragments, both sources of shearing can be measured using commonly used laboratory techniques, providing a suitable quantification of DNA integrity of DNA for sequencing with long-read technologies.

## INTRODUCTION

DNA is generally considered a relatively stable analyte as it is highly resistant to physical shear at the lengths required for PCR and short-read next generation sequencing (NGS). Protection against chemical shearing is a relatively straightforward procedure using standard methods and commonly used quality parameters (Morente et al., 2006; Lou et al., 2014; Robb et al., 2014; Ellervik and Vaught, 2015). This relative ease of handling means that there has been little need for researchers to evaluate the mechanisms of DNA fragmentation at length scales beyond a few thousand base pairs for throughout the post-genomic era (Anchordoquy and Molina, 2007; Mulcahy et al., 2016).

With the rising importance of long-read sequencing technologies and mapping technologies utilising linked reads (Schadt *et al.*, 2010; Goodwin *et al.*, 2016) shearing limits researchers from using the full capacity of emerging technologies where even megabase lengths can be sequenced in a single read (Jain et al., 2018). A better understanding of DNA structural integrity and fragmentation is therefore necessary to improve quality management practices for high-molecular-weight DNA (HMW-DNA) (English *et al.*, 2012; Malentacchi *et al.*, 2015; Goodwin *et al.*, 2016).

For short-read sequencing purposes purity, yield and amplification success are the standard measurements for DNA sample quality (Bustin *et al.*, 2009; Le Page *et al.*, 2013; Mathay *et al.*, 2016; Mulcahy *et al.*, 2016; Office of Biorepositories and Biospecimen Research *et al.*, n.d.), making it hard to estimate if DNA in samples are suitable for long-read sequencing without additional testing. The aforementioned measures are relevant for long-read sequencing as well but do not offer insight on the presence of HMW-DNA in the sample, which is necessary to generate long reads that are several thousand base pairs in length or longer. This study was therefore conducted in order to compile available knowledge on the process of DNA fragmentation and provide recommendations on how to best obtain DNA suitable for long-read sequencing.

Data on DNA integrity is yet to sparse to directly untangle the complex interdependencies between potential pre-analytical variables of relevance contributing to fragmentation (Robb *et al.*, 2014; Malentacchi *et al.*, 2016a). Therefore, this work takes an approach where experimental data is assessed and theories are collected from a wide variety of fields affected by DNA fragmentation. By merging data from the below described subjects we are able to present a model explaining the key contributing factors to DNA fragmentation when preparing biospecimen for high throughput genetics research:

- Early research on the physical properties of DNA conducted in the years after identifying the structure of the DNA helix (Bowman and Davidson, 1972; Adam and Zimm, 1977; Dancis, 1978).
- Investigations into the chemical properties and degradation of DNA in vitro/in vivo (Lindahl, 1993) that ultimately led to the discovery of cellular repair mechanisms of DNA and the Nobel prize for these discoveries.
- Archaeological research investigating the decay of DNA over long periods of time and the decay products generated during the process. Indicating that the initial rate of decay is substantially higher than can be explained by depurination (Sawyer *et al.*, 2012) but that long term fragmentation follow a 1^st^ order reaction where depurination is a key driving factor (Overballe-Petersen *et al.*, 2012) which for genomic DNA is close to what we can expect from depurination at pH 7.5 (Allentoft *et al.*, 2012).
- Forensic research provides a baseline for DNA fragmentation patterns in different tissues under conditions where no stabilization of samples is provided (Bär et al., 1988; Johnson and Ferris, 2002).
- The stability of plasmids, linearized plasmids and genomic DNA for pharmaceutical applications (Evans *et al.*, 2000). In particular, it should be noted that a similar survey to determine pharmacologically relevant factors of DNA degradation has been published (Rossmanith *et al.*, 2011). Under these conditions DNA degradation due to depurination was estimated to be the dominant factor for degradation of samples.
- Polymer science and physical chemistry studying the behaviour of polymers in turbulence (Vanapalli *et al.*, 2006), and simulation of electrostatic interactions *in silico* explaining the influence of folding and protection of DNA under high ionic-strength conditions (Cinque *et al.*, 2010; Jiang *et al.*, 2015).

A large number of potentially relevant chemical mechanisms for degradation have been identified in pharmacological research studying the degradation of DNA-based pharmaceuticals. For the purposes of DNA extraction using aqueous solutions, oxidation and depurination are estimated to be the relevant factors (Pogocki and Schöneich, 2000; Rossmanith et al., 2011). DNA sequencing of archaeological findings support these conclusions as they indicate that even DNA retrieved in conditions far from optimal show a degradation pattern consistent with depurination being the main cause of fragmentation (Allentoft *et al.*, 2012). By adjusting for this baseline of degradation other, more situational, factors for degradation such as the influence of specific DNA extraction techniques can be identified and used to generate a minimal viable explanation for DNA fragmentation under laboratory conditions.

## A MINIMAL EXPLANATION OF DNA FRAGMENTATION

When tracking DNA fragmentation in a laboratory environment the degradation process of DNA is usually considered to start at the start of the onset of the warm ischemia time for tissue collection or at the time of blood draw (or equivalent) when collecting fluids (Johnson and Ferris, 2002; Betsou *et al.*, 2010). At this time cells react to changes in the environment surrounding them, triggering processes that may cause the degradation of samples or otherwise influence experimental results. In our survey we separated the degradation into chemical shearing and physical shearing to better highlight the mechanisms of degradation where especially the physical shearing has a significantly larger influence on DNA integrity when preparing samples for long-read sequencing.

### Chemical shearing

The disruption of the cellular environment triggers a number of processes leading to the degradation of DNA by oxidizing attacks and degradation by nucleases, unless preservation measures are taken (Elmore, 2007; Rossmanith *et al.*, 2011). This process can be halted through freezing or drying of samples. It can also be prevented by the addition of preservatives, in the form of chelating agents which bind metal ions and Mg^2+^ ions, thereby preventing metal ions from catalysing the generation hydroxyl radicals and the activity of Mg^2+^ - dependent endonucleases which otherwise will attack the DNA (Lahiri and Schnabel, 1993; Rossmanith *et al.*, 2011). Chemical degradation by these pathways is the immediate threat to DNA integrity and may rapidly produce a smear of short fragments. The influence of this chemical shearing may not always be immediately apparent as “nicking” (the introduction of single strand breaks in double stranded DNA) makes the DNA highly vulnerable to future physical shearing but does not immediately fragment the DNA, making it hard for researchers to immediately recognise its influence on DNA integrity. This issue is however easily dealt with by the addition of buffers containing chelators, or a combination of EDTA + ethanol, which are highly effective in preventing degradation by oxidizing attacks (Evans *et al.*, 2000). It should, however, be noted that EDTA on its own actually catalyses the formation of hydroxyl radicals and therefore increases degradation rates unless combined with a radical scavenger (Evans *et al.*, 2000).

The other major cause of chemical shearing of DNA is depurination followed by β-elimination which is an acid-catalysed first order reaction (An *et al.*, 2014). As the reaction is driven by protonation it is impossible to completely prevent it in aqueous solutions and the complete dehydration of DNA requires encapsulation to protect the sample from atmospheric water as the DNA molecule binds water in its minor and major groove (Bonnet *et al.*, 2010). Storing samples at pH 8 (Brunstein, 2015), reducing molecular mobility, and temperature can reduce the reaction rate to levels sufficiently low to ensure that DNA maintain integrity for most long-read purposes for centuries (An et al., 2014). Apurinic sites created by depurination or depyrimidination are sensitive to β-elimination which breaks the DNA and is catalysed by primary amines (Lindahl and Andersson, 1972; Povirk and Steighner, 1989). Given the relatively high rate of β-elimination compared to depurination even in the absence of primary amines usage of Tris buffer or other compounds with primary amines should have a negligible impact on DNA integrity. Likewise the low rate of depyrimidination compared to depurination means that strand breakage due to chemical shearing is regulated by the rate of depurination in the sample if appropriate extraction methods are being used (Overballe-Petersen et al., 2012; Rossmanith et al., 2011).

### Physical stress

Susceptibility to physical stress is directly dependent on the length and folding of DNA molecules (Anchordoquy and Molina, 2007; Cinque *et al.*, 2010; Lengsfeld *et al.*, 2011). During extraction DNA is subjected to physical forces generating a peak abundance of 5-35 kb under common laboratory conditions (Malentacchi *et al.*, 2013, 2016b). Laminar flows have for a long time been seen as the main culprit behind DNA fragmentation (Dancis, 1978), but chemists specialising in the study of polymers in turbulent flow have provided convincing evidence indicating that turbulence dominates the influence of laminar flows under most conditions (Vanapalli *et al.*, 2006). These conclusions are further supported by the teams of Anchordoquy (Lengsfeld *et al.*, 2011) and Baigl (Cinque *et al.*, 2010) who have investigated the factors influencing the degradation of DNA for pharmaceutical applications (Anchordoquy and Molina, 2007).

These findings on physical shearing simplifies the work to minimise the influence of physical shear in DNA processing, as it means that most DNA fragmentation occurs when the contour length of DNA exceeds the Kolmogorov length of a turbulent flow. The Kolmogorov length scale defines the smallest possible eddies occurring in a system before they dissipate into energy. To understand this connection between quantum mechanics and biology it is necessary to know that all turbulence can be modelled as consisting of small vortices that in turn consist of smaller vortices. At a certain threshold, it is no longer possible to divide vortices into smaller ones, and at the Kolmogorov length scale these distances are so small that the spatial movement of molecules turns into vibrations (*i.e.* heat). Calculations by Vanapalli *et al.* have proven that the pull of a DNA molecule stuck in two vortices at the same time is sufficient to break the strand in half and that this is the primary cause of fragmentation under most hydrodynamic conditions (Vanapalli *et al.*, 2006). During sample processing fragmentation occurs over time until all DNA molecules have been degraded to a length where all fragments are short enough to remain within a single eddy, which is consistent with experimental results with commonly used laboratory equipment, such as vortex mixers and pipettes (Yoo *et al.*, 2011), and results suggest that the breakage rate is a function of the shear rate rather than the shear stress (Bowman and Davidson, 1972; Meacle *et al.*, 2007).

### Freezing and long-term storage

Finally, DNA can be fragmented by freezing, thawing, and long-term storage. Repeated freeze-thaw cycles are the most significant source of extensive damage but continued chemical reactions while the samples are frozen also play a role. Researchers frequently assume that nothing happens to DNA inside a frozen sample (Hubel *et al.*, 2014), but in reality both chemical processes and physical movement occur even at low temperatures. Between −39 and −50 °C, the DNA molecule becomes rigid and brittle as it enters a glassy state (Norberg and Nilsson, 1996) while protons in the water remain mobile even at temperatures close −268 °C (Benton *et al.*, 2016). There is a lack of long term studies at −80 °C or −196 °C, and the complex behaviour of water at low temperatures makes it impossible to estimate at what temperatures chemical activity generated by protons ceases to occur. Using the Arrhenius equation to estimate an upper bound of activity we can, however, estimate that a strand break occur at a rate of approximately once every 5 seconds in a 3.2 billion base pair genome at 25 °C and pH 8. Storage in a −80 °C freezer reduces the rate to less than once per 10 000 years (An et al., 2014).

During freezing and thawing two other factors may contribute to the fragmentation of DNA (Anchordoquy *et al.*, 2001; Shao *et al.*, 2012). As a sample cools beneath the freezing temperature of fluids, it will undergo nucleation where water molecules (or other fluids) start assembling into crystals. Solutes are not incorporated into the solid ice crystals, resulting in the sample separating into solid components and unfrozen liquid components with high concentration of solutes. Creating an environment where two sources of DNA shearing has been proposed (Anchordoquy *et al.*, 2001; Brunstein, 2015):

- As ice crystals form, solutes are crowded into gaps with increasingly high concentrations of solutes, thereby drastically increasing the potential for oxidising attacks and protonation of the DNA chain, leading to chemical shearing of the DNA polymer.
- As ice crystals form, DNA polymers are partially embedded within the ice crystals. The mechanical stress could thereby lead to the breakage of DNA molecules with a higher likelihood of breakage occurring in longer polymers.

Based on laboratory experiments, it is evident that chemical shearing is present under certain circumstances (Davis et al., 2000) but can be completely avoided as highly purified samples of short DNA fragments (<2 kb) have remained intact during repeated freeze-thawing in several studies (Ross *et al.*, 1990; Thornton *et al.*, 2005; Integrated DNA Technologies, 2014; Rossmanith *et al.*, 2011). Under certain conditions shearing also appears to be length dependent as demonstrated by Rossmanith *et al.* as short amplicons (274 bp) remained intact during experiments while the same amplicons when located within longer genomic sequences from *Salmonella typhimurium* (4 857 kb) and *Listeria monocytognes* (1 160 kb) were sheared during repeated freeze-thaw cycles (Rossmanith *et al.*, 2011).

Research in cryopreservation of oocytes, embryos and sperm (Amidi *et al.*, 2016; Kopeika *et al.*, 2015) suggest that shearing of DNA during freeze-thaw can be reduced by the addition of antioxidants, indicating that oxidation of DNA by radicals may, at least *in vivo*, be a significant cause of DNA fragmentation. Such conclusions must however be considered highly tentative as there are substantial differences in the chemical environment inside the cell compared to the *in vitro* conditions covered by this study. The safeguards necessary to avoid physical shearing by ice crystal formation are also poorly understood. The addition of glycerol (Calcott and Gargett, 1981; Schaudien *et al.*, 2007) is known to protect DNA during freeze-thaw cycles but we have been unable to determine why samples seemingly subjected to physical shearing may converge to lengths anywhere from below 1000 bp and 25 000 bp in different studies (Röder et al., 2010; Ross et al., 1990; Rossmanith et al., 2011; Schaudien et al., 2007; Shao et al., 2012).

## DISCUSSION

Long-read sequencing enables studies of complex genomic events, e.g. structural re-arrangements, copy number variations, repeat expansions, etc. Hence, it is obvious that the long-read technologies, such as PacBio, 10x Genomics and Oxford Nanopore will be increasingly popular. This constitutes a challenge for laboratories and biobanks, as best practice routines for storage of biological samples and DNA extractions need to be adjusted in order to allow delivery of HMW-DNA suitable for long-read sequencing (Malentacchi *et al.*, 2016a). Currently, the Oxford Nanopore MinION, GridION and PromethION are able to sequence reads up to 1 Mbp in length (Oxford Nanopore Technologies, 2017). The optimal results obtained by 10x Chromium systems are obtained if the sample contains a high number of molecules above 50 kbp, and up to a few hundred kilobases, and the PacBio systems can produce reads up to 60-70 kbp (Bickhart et al., 2017; Lok et al., 2017; Paajanen et al., 2017). These numbers will most likely rise due to technology development, and processing facilities and biobanks should be ready to provide researchers with DNA samples of adequate quality and quantity.

Current biobanking practices provide DNA with a high yield and purity when tested, but suffer from a high level of variability in regards to DNA integrity (Malentacchi et al., 2015, 2013). Improved protocols and subtle alterations of protocols can, however, provide significant improvements. The SPIDIA-ring trials provide one such valuable example as they show that biobanks are able to rapidly improve their operations when provided with sufficient guidance (Malentacchi *et al.*, 2016a). It is therefore important that an acceptable definition of DNA fragmentation and robust units of measurement are made available to the research community to improve the availability of HMW-DNA for researchers. In this regard the SPIDIA project (Malentacchi *et al.*, 2013) and the Global Genome Biodiversity Network (GGBN) (Mulcahy *et al.*, 2016) have provided pioneering work on the subject on how to objectively evaluate and compare DNA fragmentation in samples using gel-electrophoresis and a *Hin*dIII ladder. Both projects propose a “greater than X kb” approach where the percentage of DNA above a certain “DNA threshold” is measured and used as an indicator of DNA integrity. The SPIDIA project used the 4.36 kb size marker as its X-value and supplemented it with the measured peak intensity to provide an assessment of DNA integrity (Ciniselli *et al.*, 2015). The GGBN on the other hand decided to forego the usage of a peak intensity and instead proposed the X value to be dependent on the intended utilization of samples but with the 9.416 bp size marker as a default as it is substantially longer than standard HTS reads and would be an appropriate minimum for long-read sequencing (Mulcahy *et al.*, 2016).

The influence of physical shear stress and chemical degradation are, however, substantially different from each other. We therefore believe that two separate measures of DNA fragmentation are necessary to help researchers to evaluate the integrity of their DNA in an adequate manner. Thereby providing an assessment of both the random chemical shearing currently measured and length-dependent physical shearing which limits the availability of samples suitable for long-read sequencing.

**The characteristic fragment length (CFL):**The most abundant fragment length of the sample. In gel-electrophoresis (both conventional and capillary), this value corresponds to the peak intensity of the signal.

**The smear ratio (SR)**: The proportion of DNA with a fragment length of less than 500 bp. This value can be easily calculated using the <u>imageJ</u> open source software to integrate over the signal intensity curve as described by Mulcahy *et al.* (Mulcahy *et al.*, 2016).

The characteristic fragment length is routinely reported when DNA integrity is assessed using gel-electrophoresis. Based on the conclusions drawn in this paper it can be deemed suitable as a direct but conservative estimate of how physical shear stress limits the DNA fragment length in a sample.

The smear ratio provides an estimate of the proportion of DNA that has been shredded to short fragment lengths due to freeze-thaw shearing or chemical degradation. From a technical perspective the work to produce a “less than X kb” measurement is identical to the “greater than X kb” proposed by GGBN, but it provides researchers with a more informative measurement. Samples with highly preserved content will have a lopsided distribution around the characteristic fragment length with a long tail of shorter DNA fragments. Changes in the maximum fragment length by physical shear stress will easily be detected by a shift in peak fragment length and a “less than X kb” measurement provides a corresponding estimate of the amount of DNA being sheared to very short fragments due to chemical degradation and/or freeze-thaw.

These two measurements therefore provide laboratories with the means to monitor changes in processing and allow end users to assess the potential of obtaining sufficiently HMW DNA from a resource. For advanced users and quality management it is, however, still attractive to also make the entire fragment distribution plot available as there is often a “plateau” of HMW-DNA preceding the peak centred around the characteristic fragment length (figure 1). Based on the experience of National Genomics Infrastructure - Sweden, the following instrumentation can provide researchers with this type of information providing both the characteristic fragment length and smear ratio in a single analysis (figure 1): Agilent Bioanalyser (Agilent Technologies), Fragment Analyzer (Advanced Analytical) and Femto Pulse (Advanced Analytical) with the Femto pulse being preferred for measuring DNA fragments longer than 10 kb at the National Genomics Infrastructure.

**Figure 1.**
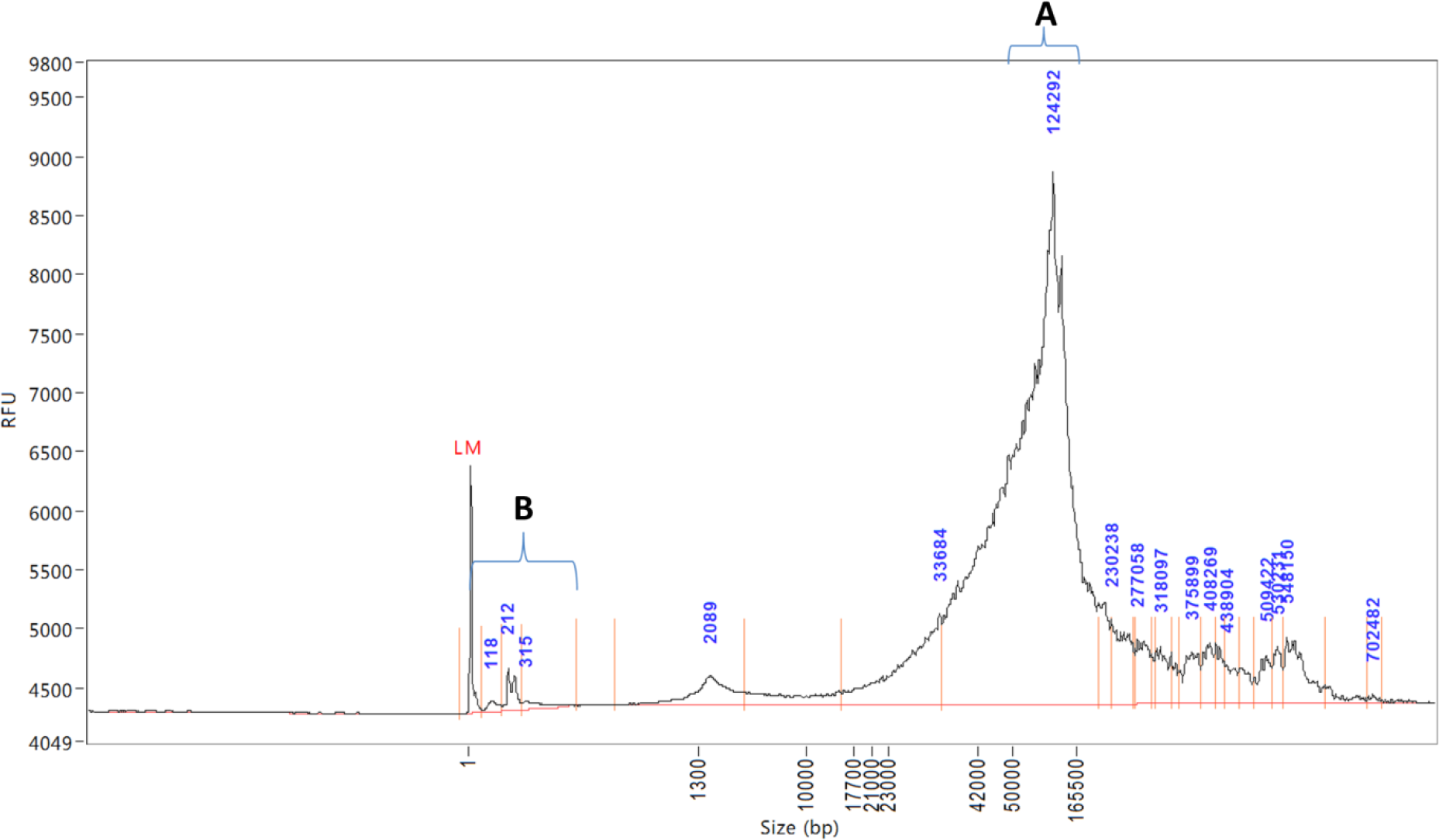
example of high quality DNA with minimal smear (section B) and a characteristic fragment length of 129 292 bp as measured by the Femtopulse (Advanced Analytical). High quality samples often contain a plateau on the right hand side of the characteristic fragment length, meaning that the CFL should not be interpreted as a measure of the longest fragment length available for sequencing. To provide longer reads suitable for long-read sequencing or linked-read technology, the DNA extraction process can be altered to reduce physical shearing. With continuous monitoring it is also possible to quickly any inadvertent changes influencing DNA integrity as chemical shearing an increased smear ratio or a drop in characteristic fragment length is immediately measured using the above described technique.

The first megabase plus read was recently announced using the MinION technology from Oxford Nanopore (Oxford Nanopore Technologies, 2017) and fragments longer than two megabases can be reconstructured from fragments separated during sequencing (Payne et al., 2018). At such long-read lengths, currently standard methods require modifications and gentle DNA extraction methods such as phenol-chloroform extraction (for reference the Rad003 protocol used by Loman lab to extract DNA for generating the megabase-length read is available at http://lab.loman.net/protocols/). The shearing of DNA caused by the extraction step itself is, at least in high-throughput laboratories such as biobanks, likely to generate fragments with a characteristic length of more than 50 kb, while turbulent forces during other parts of the process restrict the fragment length to 5-35 kb (Shao *et al.*, 2012; Malentacchi *et al.*, 2015). Among commonly used extraction methods, precipitation based DNA extraction generate the highest molecular weight of DNA (Malentacchi *et al.*, 2015) but variations in fragment length (Shao *et al.*, 2012) and other quality parameters mean that kit-by-kit comparisons will be necessary for the selection of suitable best practice procedures for DNA extraction depending on the intended usage of the samples.

Nanopore sequencing provides the longest continuous reads in sequencing and there is significant optimisms with researchers suggesting that read-lengths are dependent on the input fragment length (Jain et al., 2018; Magi et al., 2017). Careful preservation of HMW-DNA has also enabled feats such as the complete assembly of the human major histocompatibility complex (MHC) in a single contig (Jain et al., 2018) showing the significant value of carefully preserved DNA. It should however be noted that there is little published information regarding the influence of contaminants on the read length of DNA which is a topic outside the scope of this study and there is also evidence that recorded reads may be shorter than the fragments present in the sample during adapter ligation (Payne et al., 2018). If this is the result of breakage during the sequencing itself, a technical artefact from data processing or the result of inhibitors limiting the effective read length is currently unknown and requires further research. It does however indicate that even if Nanopore sequencing theoretically is limited by the input fragment length, there is still a need to assess DNA integrity for quality management purposes rather than rely on the read length distribution generated by sequencing with the nanopore.

## CONCLUSIONS

Based on the model presented above, laboratories are advised to focus on three areas if they wish to increase the integrity of extracted DNA samples (table 1). During the retrieval of samples it is important to quickly stabilise the sample by preventing enzymatic degradation and generation of free radicals within the sample. After the cells have been lysed it is important to handle the sample with outmost care to minimise physical shearing (gentle mixing, slow pipetting, using wide-bore pipette tips, etc) and then store DNA at low temperature or in a completely dehydrated state to avoid depurination.

**Table 1.** key parameters for identifying causes of DNA fragmentation and how to reduce it.

|  | Source | Time scale | Mechanism | Threat profile | Preventive measure | Damage pattern |
| --- | --- | --- | --- | --- | --- | --- |
| Sample collection | Enzymatic degradation | Immediate | Naturally occurring DNAses attack the DNA polymer | An immediate threat to the integrity of DNA unless enzymes are inhibited or destroyed. | Addition of Proteinase degrading the protein or chelators binding metal ions functioning as co-factors. | DNA fragmentation with specific cleavage sites |
|  | Oxidation | Seconds to years | Metal ions catalysing the production of hydroxyl radicals | An immediate threat to the integrity of DNA unless chelating agents and/or a radical scavenger is added. | Addition of chelators or EDTA + Ethanol | Smearing |
| Processing | Turbulence | Immediate | Physical shearing | Poorly optimised machinery and careless handling of sample will cause physical shearing of the DNA. | Freezing, dehydration or maintaining a high P/E-value | Characteristic fragment length |
|  | Depurination | Weeks to years | Protonation of purines lead to depurination followed by beta elimination | A slow reaction that may cause issues under acidic conditions or long term storage at room temperature. |  | Smearing |
| Storage | Freeze-thaw | Immediate | Physical shearing | The damage mechanisms of freeze-thawing DNA are still poorly understood *in vitro* but damage is proportional to the number of freeze-thaw cycles the sample has undergone. Flash freezing or the addition of a compound allowing samples to vitrify rather than form ice crystals seem effective. | Addition of a protective agent such as glycerol or flash freezing to low temperatures in order to prevent large ice crystals from forming. | Characteristic fragment length |
|  | Freeze-thaw | Immediate | Crowding of solutes leading to hydrolysis and/or oxidation | The damage mechanisms of freeze-thawing DNA are still poorly understood *in vitro* but damage is proportional to the number of freeze-thaw cycles the sample has undergone. Flash freezing or the addition of a compound allowing samples to vitrify rather than form ice crystals seem effective as solutes are not crowded into the vicinity of the DNA polymer. Minimising the number of freeze-thaw cycles is also an obvious solution. | Maintaining a high purity of extracted DNA and possibly addition of protective agents. | Smearing |
|  | Depurination | Weeks to years | Protonation of purines leading to depurination followed by beta elimination | A slow reaction that may cause issues under acidic conditions or long term storage at high temperature or with insufficient removal of water | Low temperature storage or complete dehydration. | Smearing |

Physical degradation by shear stress is mainly caused by microscale turbulence. The turbulence shears DNA until the fragments reach a characteristic peak length. Physical forces are strongest in the centre of the DNA polymer but nicks and random events may cause strand breaks elsewhere. This generates a characteristic peak length roughly corresponding to half the length of the smallest fragment length prone to degradation in the system. When measured by gel electrophoresis this generates a characteristic pattern with a peak intensity of fluorescence followed by a long tail smear below the peak and a shorter upstream section of fragments that, due to random/chemical effects, reached a length protected from turbulence but still longer than half the critical length.

Freeze-thaw events may contribute significant damage to DNA but is highly dependent on parameters such as polymer size, ion strength, rate of temperature drops, presence of nicks etc. Currently, there is no generalized theory to predict fragmentation but preservation with means such as storage in 50 % glycerol has been shown to prevent shearing. Furthermore, it seems that variations in upstream protocols or the concentration of DNA in samples have a profound impact on the vulnerability to freeze-thaw cycles as evidenced by in-house testing not identifying any fragmentation of samples exposed to repeated freeze-thaw cycles.

To support further work on maintaining DNA integrity it is important that researchers report relevant data on the integrity of DNA. DNA integrity data is often omitted or reported using inadequate measurements such as electrophoresis with *Hind*III digested λ DNA, which does not provide relevant information above 23 kb. Current literature on maintaining longer DNA for high throughput applications is therefore limited and this study provides a foundation for further work in quality management and process optimisation for the production of HMW-DNA for sequencing.

## ACKNOWLEDGEMENT

The authors would like to thank Dr Jaan Hong for his help with drawing our blood as well as PhD candidate Ida Höijer and Dr Oskar Karlsson for their help in the laboratory that have allowed us to run confirmatory tests.

## Funding

This work was financed by the BBMRI-LPC and the B3Africa projects. BBMRI-LPC is supported by the European Community’s 7th Framework Programme (FP7/20072013) grant agreement no. 313010, B3Africa is supported by the European Union s Horizon 2020 research and innovation programme under grant agreement No 654404. Work performed at Uppsala Genome Center – NGI Uppsala has been funded by RFI – VR and SciLifeLab Sweden.

